# PSGL-1 excludes HIV Env from virion surface through spatial hindrance involving structural folding of the decameric repeats (DR)

**DOI:** 10.1101/2024.12.28.630612

**Authors:** Sameer Tiwari, Bryan M. Delfing, Yang Han, Christopher Lockhart, Amrita Haikerwal, Abdul A. Waheed, Eric O. Freed, M. Saleet Jafri, Dmitri Klimov, Yuntao Wu

**Author notes:** Correspondence and requests for materials should be addressed to Y.W. or D.K. **Email:**.

## Abstract

P-selectin glycoprotein ligand-1 (PSGL-1), a mucin-like surface glycoprotein, is primarily expressed on lymphoid and myeloid cells. PSGL-1 has recently been identified as an HIV restriction factor, blocking HIV infectivity mainly through virion incorporation that sterically hinders virion attachment to target cells. PSGL-1 also inhibits HIV Env incorporation into virions. However, the molecular mechanisms of PSGL-1-mediated Env exclusion remained unclear. Here, we investigated the role of PSGL-1’s extracellular (EC) and intracellular (IC) domains in Env exclusion. We demonstrate that both EC and IC are important for Env exclusion; when EC was deleted, PSGL-1 completely lost its ability to inhibit Env incorporation, whereas when IC was deleted, PSGL-1 partially lost this activity. In addition, when the decameric repeats (DR) were deleted from EC, PSGL-1 also lost its ability to inhibit Env incorporation. Sequential DR deletion mutagenesis further demonstrated that a minimum of 9 DRs is necessary for Env exclusion. Molecular modeling of the DR structure revealed that PSGL-1 mutants with 7 or fewer DRs pose as an extended “rod-like” structure, whereas those with 9 or more DRs collapse into a “coil-like” structure that spatially excludes Env. Our studies suggest a model in which Env exclusion involves Gag-mediated PSGL-1 targeting to the virion assembly site where DR-mediated spatial exclusion blocks Env incorporation.

## Introduction

P-selectin glycoprotein ligand-1 (PSGL-1), or CD162, is a 120-kDa surface glycoprotein that binds to the selectin family of proteins, including P-, E-, and L-selectin, with the highest affinity for P-selectin ^1–3^. PSGL-1 is primarily expressed on the surface of lymphoid and myeloid cells ^1,4,5^ and functions to recruit lymphocytes into inflamed tissues ^6–9^. The recruitment process involves PSGL-1 binding to selectins expressed on the surface of endothelial cells of the blood vessel. This interaction initiates leukocyte tethering and rolling on the endothelium surface for migration into inflamed tissues ^6–9^. Structurally, PSGL-1 is characterized by a mucin-like, N-terminal extracellular domain (EC) and a C-terminal intracellular domain (IC). EC has a highly extended structure, with the extracellular portion projecting nearly 60 nm from the membrane surface ^10,11^. EC is also heavily glycosylated and is relatively rigid, comprising 14-16 tandem decameric repeats (DRs) ^12,13^. Each DR consists of a repeated sequence of 10 amino acids (-A-T/M-E-A-Q-T-T-X-P/L-A/T-) and contains multiple O-glycosylated threonines (∼30%) and prolines (∼10%) ^10,11,13^. IC contains a polybasic region that can interact indirectly with the highly basic region (HBR) of the HIV matrix protein (MA) ^14,15^. This interaction has been found to mediate PSGL-1/Gag colocalization to the site of virion assembly ^14^.

PSGL-1 restricts HIV-1 infectivity mainly through virion incorporation that sterically hinders virus attachment to target cells ^16–18^. In addition, during viral assembly, the presence of PSGL-1 in virus-producing cells diminishes the levels of envelope proteins incorporated into virions ^17^. The mechanisms of HIV Env incorporation have remained largely elusive and may involve specific or non-specific interactions of Gag with Env, or Gag and Env co-targeting ^19^. Based on the current models of Env incorporation ^19^, we hypothesized that PSGL-1 may exclude Env by blocking Env display on the plasma membrane through spatial competition resulting from the highly glycosylated extracellular domain of PSGL-1. Additionally, PSGL-1 may exclude Env incorporation through intracellular competition for MA binding, either directly or indirectly. To this end, we probed the roles of the EC and IC domains in Env exclusion through PSGL-1 deletion mutagenesis.

## Results

### Role of PSGL-1’s extracellular (EC) and intracellular (IC) domains in HIV Env exclusion

To determine the roles of PSGL-1’s extracellular (EC) and intracellular (IC) domains in Env exclusion, we selected three EC or IC deletion mutants (PSGL-1-NT, -CT, and -ΔCT) (**Fig. 1a**). The mutant DNAs were co-transfected with HIV-1(NL4-3) DNA into HEK293T cells. Virions were assembled and harvested at 48 hours, purified through 10% sucrose cushion, and analyzed with Western blotting to detect virion HIV gp41 and p24. As shown in **Fig. 1b**, full-length PSGL-1 inhibited Env incorporation ^17^, whereas deleting EC or IC led to a decrease in the inhibition of Env incorporation; in particular, deleting EC (PSGL-1-CT) led to a complete loss of the inhibition (**Fig. 1b**), while deleting IC led to a partial loss of the inhibition These results demonstrated that although both EC and IC are involved in Env exclusion, EC plays a more pronounced role.

**Figure 1.**
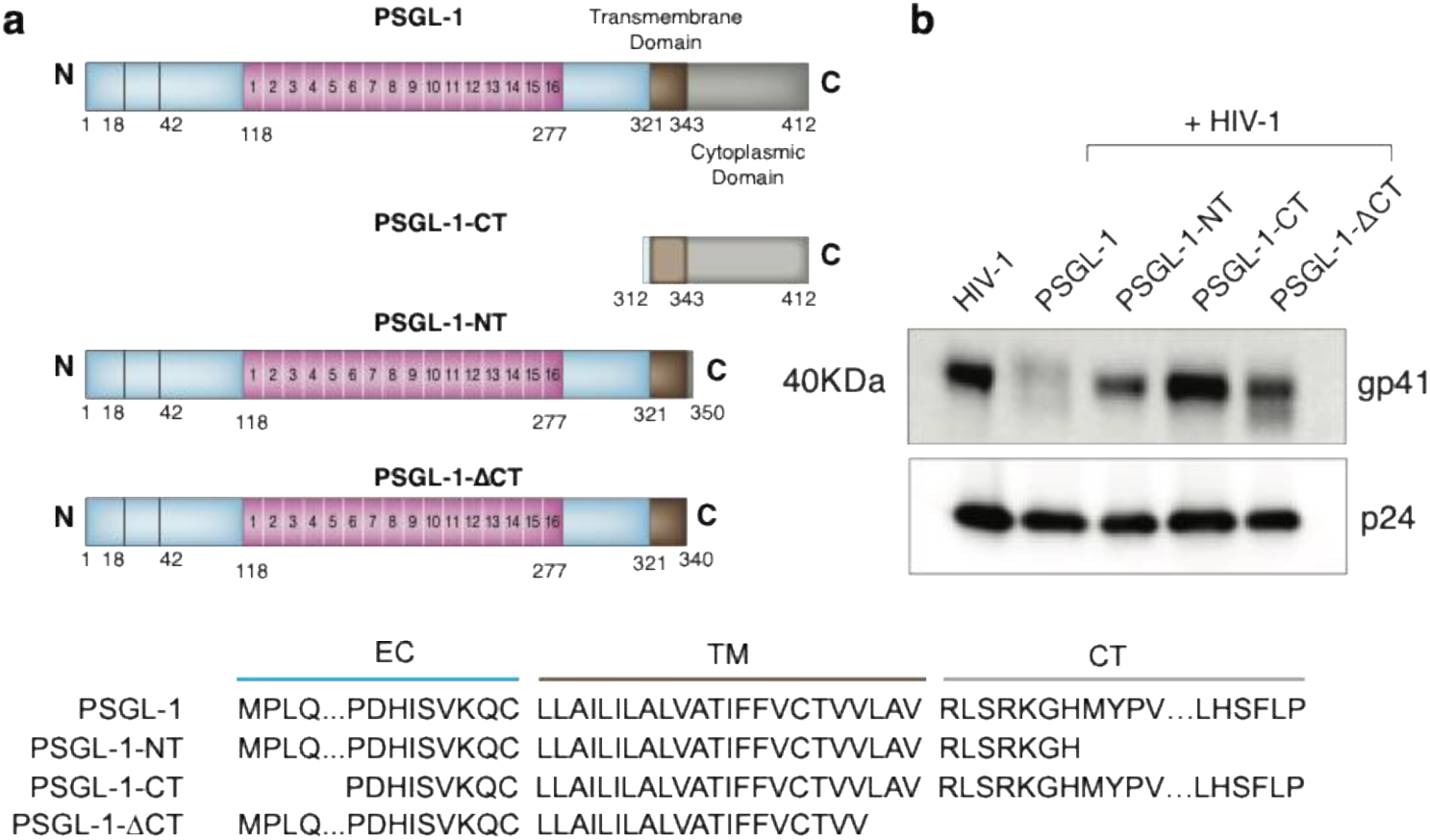
Requirement of PSGL-1’s N-terminal and C-terminal domains for blocking Env incorporation. (**a**) Schematic diagram showing PSGL-1’s EC and IC deletion mutants, PSGL-1-NT, PSGL-1ΔCT, and PSGL-1-CT. The amino acid sequence alignment of the deletion mutants shows PSGL-1’s extracellular (EC), transmembrane (TM), and C-terminal (CT) domains. (**b**) Western blot quantification of HIV gp41 levels in virions produced from cells co-transfected with full-length PSGL-1 DNA or its NC or IC deletion mutants DNA. Virion p24 was also probed and used for normalizing levels of particles. Normalized levels of gp41 are shown as fold change. The experiment was repeated three times independently.

### A minimum of 9 DRs within EC is necessary for Env exclusion

EC contains the decameric repeats (DRs), comprising 14-16 tandem decameric repeats (-A-T/M-E-A-Q-T-T-X-P/L-A/T-) ^12,13^ which are heavily glycosylated and contain multiple O-glycosylated threonines (∼30%) and prolines (∼10%) ^10,11,13^. We further tested the role of DRs in HIV Env exclusion by creating a panel of sequential DR deletion mutants carrying various numbers of DRs (**Fig. 2a**). When all 16 DRs were deleted (PSGL-1-ΔDR), PSGL-1 lost its ability to block Env incorporation (**Fig. 2b)**; partial deletion of DR to 1 to 7 DRs (PSGL-1-1DR to PSGL-1-7DR) also largely abolished PSGL-1’s ability to exclude Env (**Fig. 2b**). However, when the numbers of DRs were kept at 9 or more, PSGL-1 maintained its ability to exclude Env (**Fig. 2b,c**). These results demonstrated that a minimum of 9 DRs is necessary for Env exclusion.

**Figure 2.**
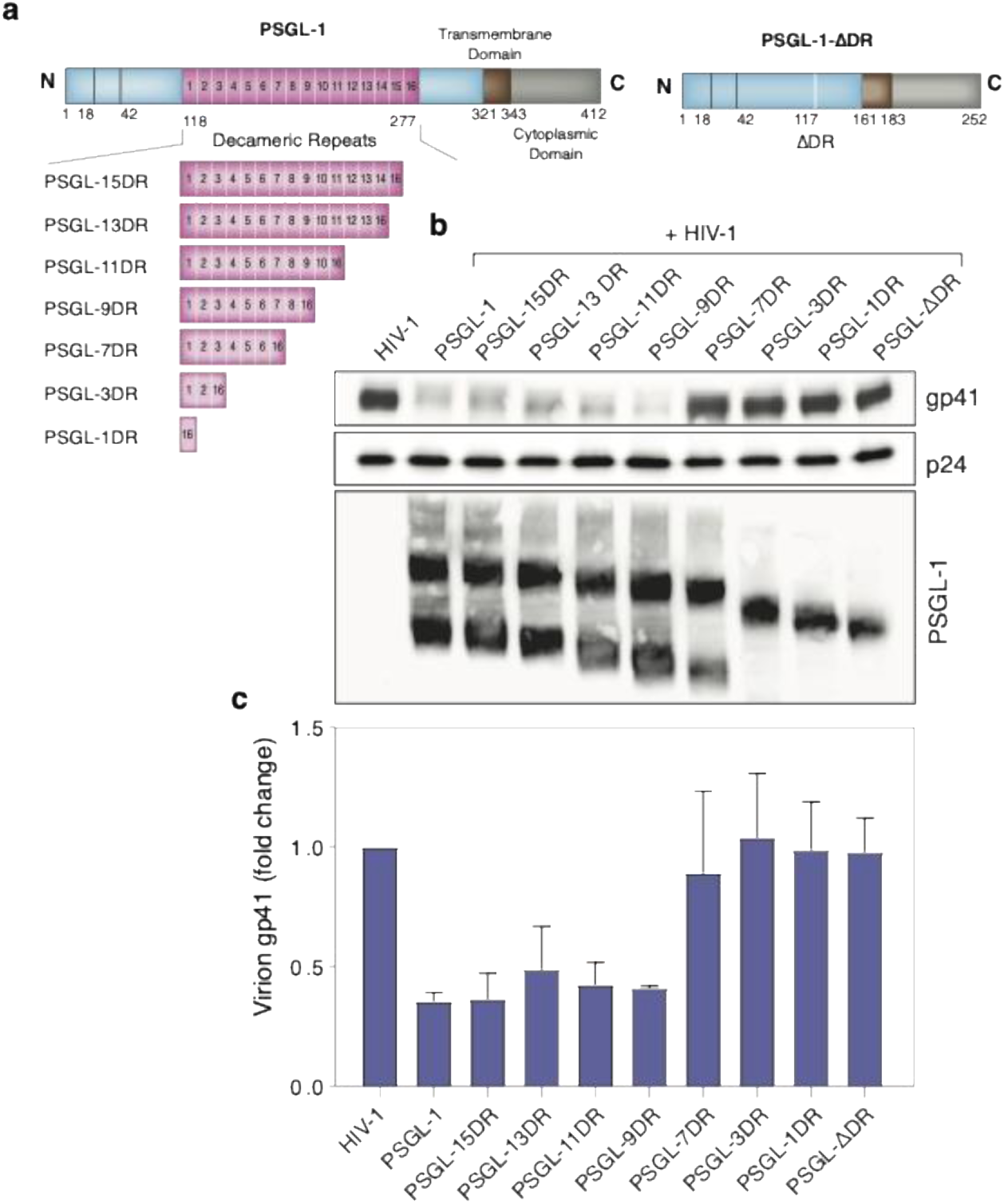
Requirement of PSGL-1’s DR domain for blocking Env incorporation. (**a**) Schematic diagram showing sequential DR deletions in PSGL-1 DR mutants. (**b**) Western blot quantification of HIV gp41 levels in virions produced from cells co-transfected with full-length PSGL-1 DNAs or its DR deletion mutants DNA. Virion p24 was also probed and used for normalizing levels of particles. Normalized levels of gp41 are shown as fold change. Levels of virion PSGL-1 were also quantified. (**c**) The experiment was repeated three times independently, and the normalized gp41 levels were quantified and are shown as the averages and standard deviations.

### Molecular modeling of the DR structure reveals two distinctive structural conformations defined by the number of DRs

To understand the molecular basis for this minimal requirement of 9 DRs for HIV Env exclusion, we used AlphaFold 3 ^20^ to predict PSGL-1 DR structures for the repeats composed of 1, 3, 5, 7, 8, 9, 10, 11, 13, 15, and 16 DRs, with or without glycans (**Tables 1 to 4**). The structures of DR repeats were characterized using three measures (**Fig. 3**). The end-to-end distance, *r*, measures the separation between the first and last Cα carbons in a given DR repeat. The radius of gyration, *R_g_*, measures the overall dimensions of the DR repeats. To compute *R_g_*, we used non-glycan heavy atoms in a DR repeat and the “measure rgyr” function in VMD (visual molecular dynamics) ^21^. To map spatial distribution of atoms in a repeat, we computed the probability distributions of its heavy atoms *P*(*r*) as a function of their distance *r* to its geometric center. A bin size of 0.5 Å was used to compute *P*(*r*). As shown in **Fig. 3**, our modeling data suggest that an increase in the number of DRs results in complex, extension-collapse structural changes in DR repeats. As evidenced in **Fig. 4** and confirmed in **Fig. 3**, DR repeats behave as “rod-like” extended objects as their length *n* increases from 1 to 7 DRs; however, when the number of DRs *n* reaches 8, the repeats start to collapse, coinciding conspicuously with the onset of HIV Env exclusion when there are more than 7 DR repeats (**Fig. 2b**). This collapsing effect continues as the number of DRs in a tandem increases until *n* = 16. The changes in DR structure suggest that rigid, extended repeats do not impart Env exclusion, which instead is apparently driven by the repeat collapse into a larger “coil-like” structure. We concluded that spatial excluded volume interactions offered by the collapsed DR repeats are the possible reason for Env exclusion.

**Table 1.**
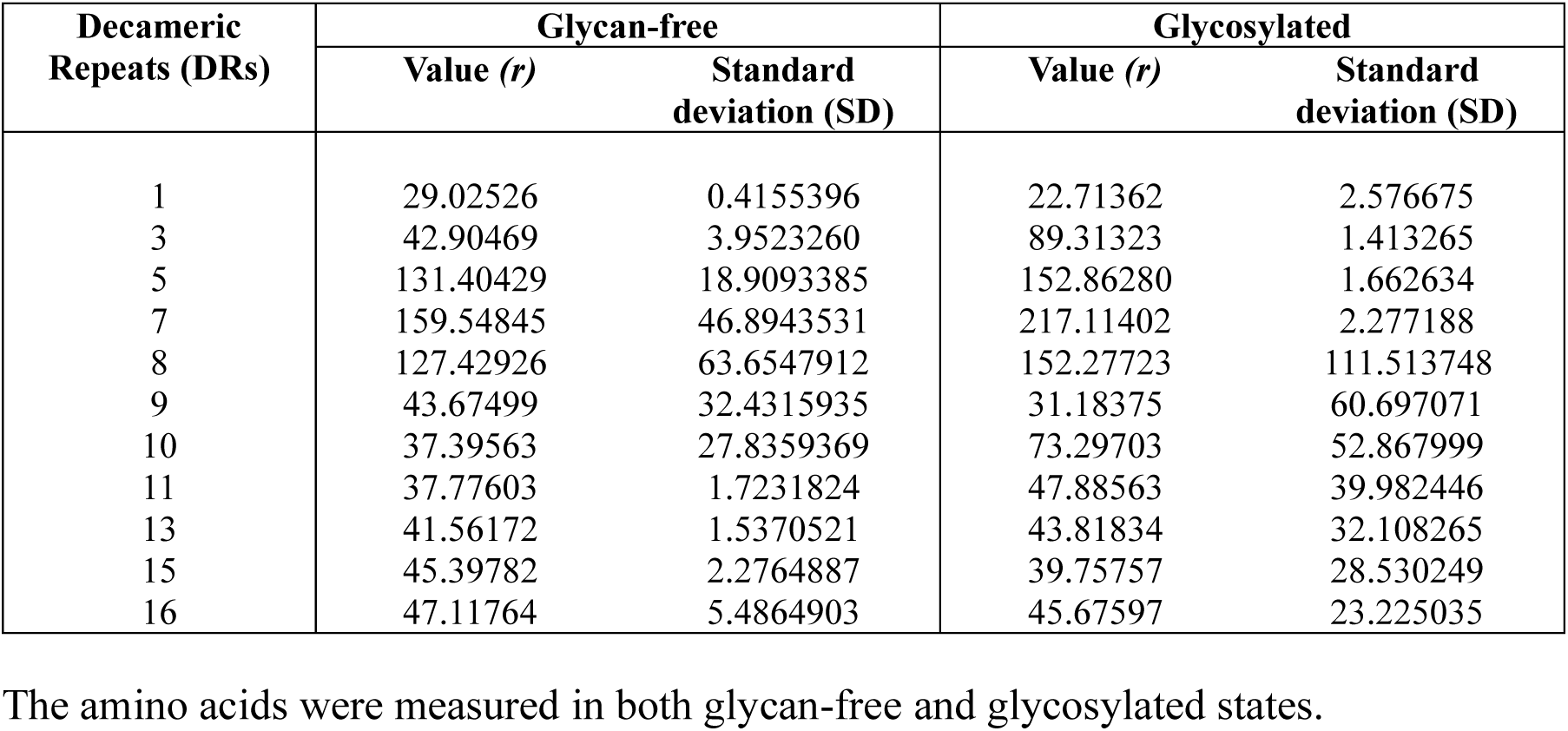
Raw data to Fig. 3a, calculation of End-to-end distance, *r*, for DR repeats as predicted by AlphaFold3.

**Table 2.** Raw data to Fig. 3b, Calculation of radius of gyration, *R_g_*, of DR repeats as predicted by AlphaFold3.

| Decameric Repeats (DRs) | Glycan-free |  | Glycosylated |  |
| --- | --- | --- | --- | --- |
| | Value ($R_g$) | Standard deviation (SD) | Value ($R_g$) | Standard deviation (SD) |
| 1 | 9.300588 | 0.1117385 | 8.085996 | 0.5950743 |
| 3 | 13.805861 | 0.8767563 | 27.427636 | 0.3522424 |
| 5 | 39.594516 | 5.8027083 | 46.005786 | 0.3305210 |
| 7 | 46.633063 | 14.4115404 | 64.440770 | 0.5583026 |
| 8 | 36.322733 | 15.7054146 | 59.479130 | 16.0506388 |
| 9 | 14.544877 | 7.9090459 | 44.699530 | 8.8746234 |
| 10 | 14.412309 | 7.0114033 | 38.590831 | 7.3997469 |
| 11 | 13.425455 | 0.1590898 | 29.222780 | 4.2318436 |
| 13 | 14.468012 | 0.3953240 | 25.487617 | 3.638325 |
| 15 | 15.318359 | 0.2209248 | 23.262024 | 2.7813683 |
| 16 | 15.954491 | 0.3254726 | 21.237016 | 2.2998281 |

The radius of gyration measures the overall dimension of a repeat. The amino acids were measured in both glycan-free and glycosylated states.

**Table 3.**
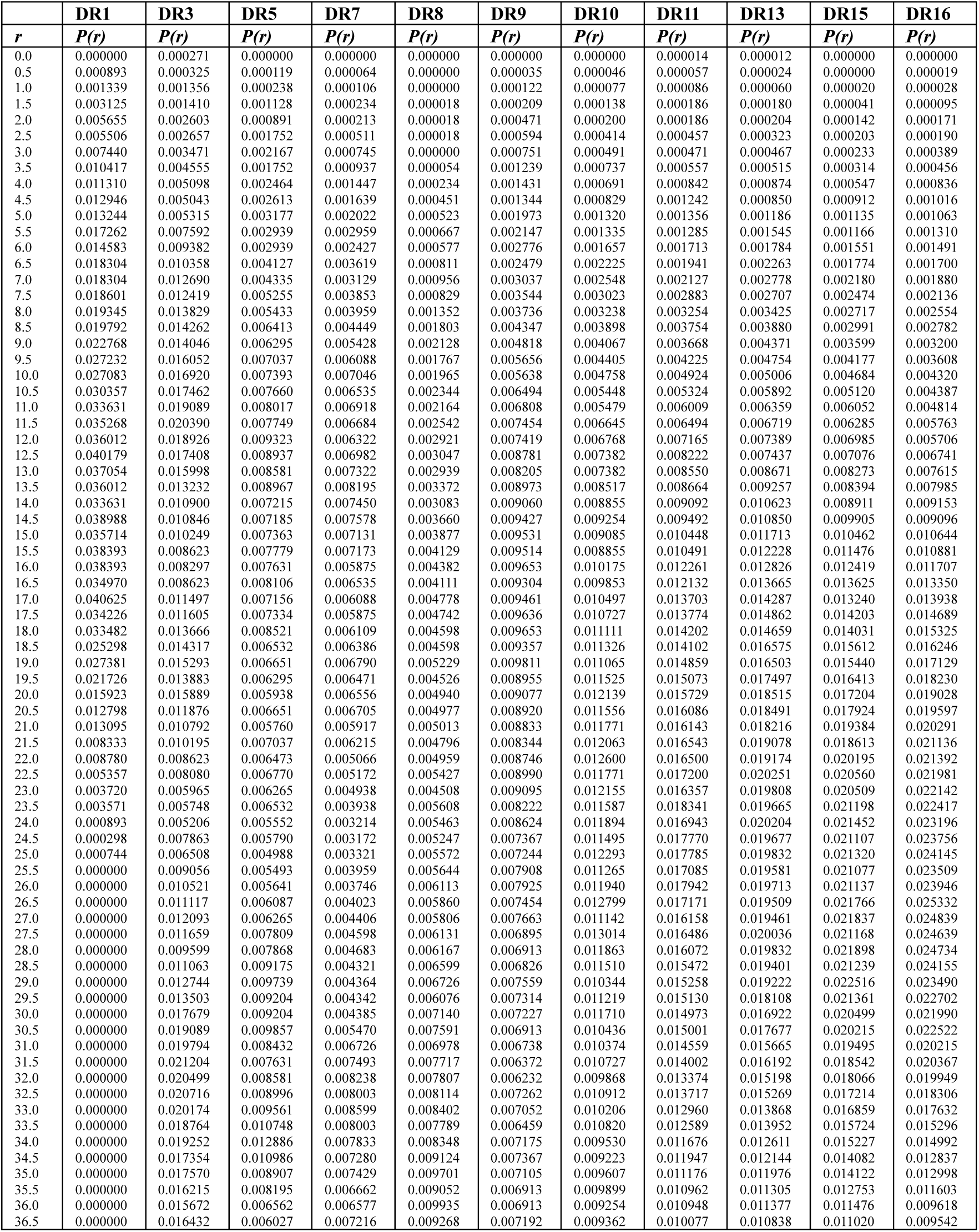
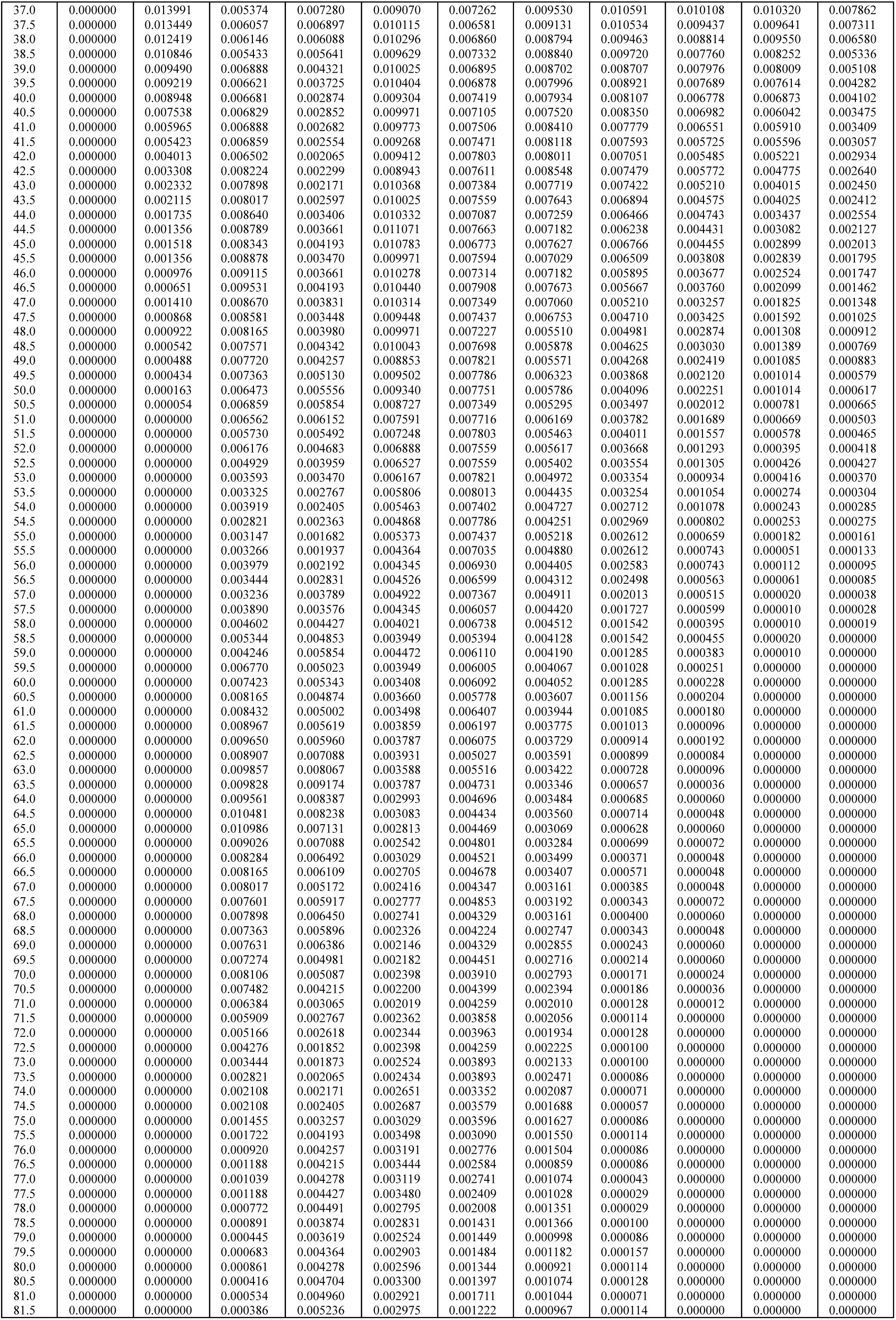
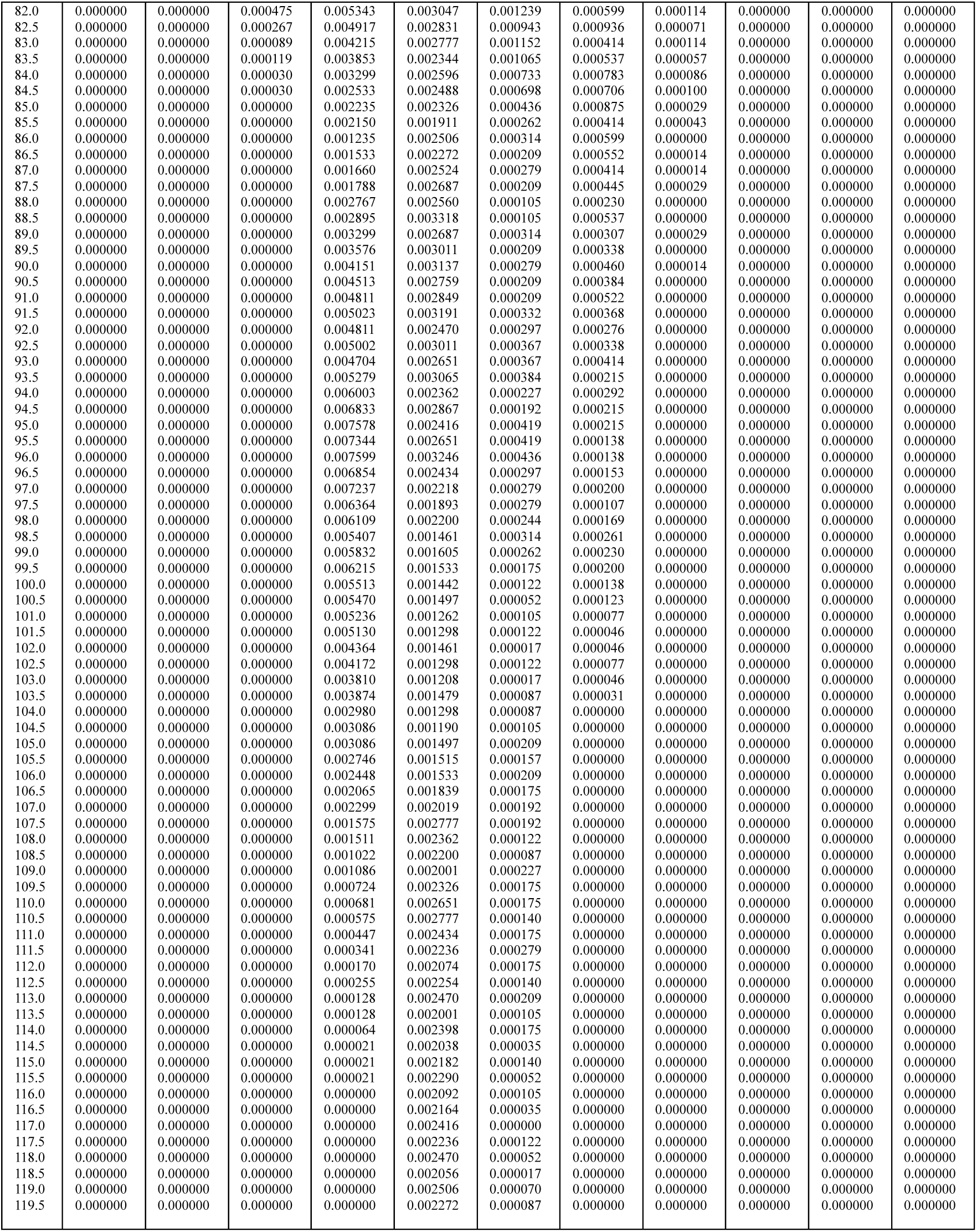
Raw data to Fig. 3c, Probability distributions *P (r)* of repeat atom distances, r, from the geometric centers of the repeats are shown for glycosylated repeats as predicted by AlphaFold3.

**Table 4.**
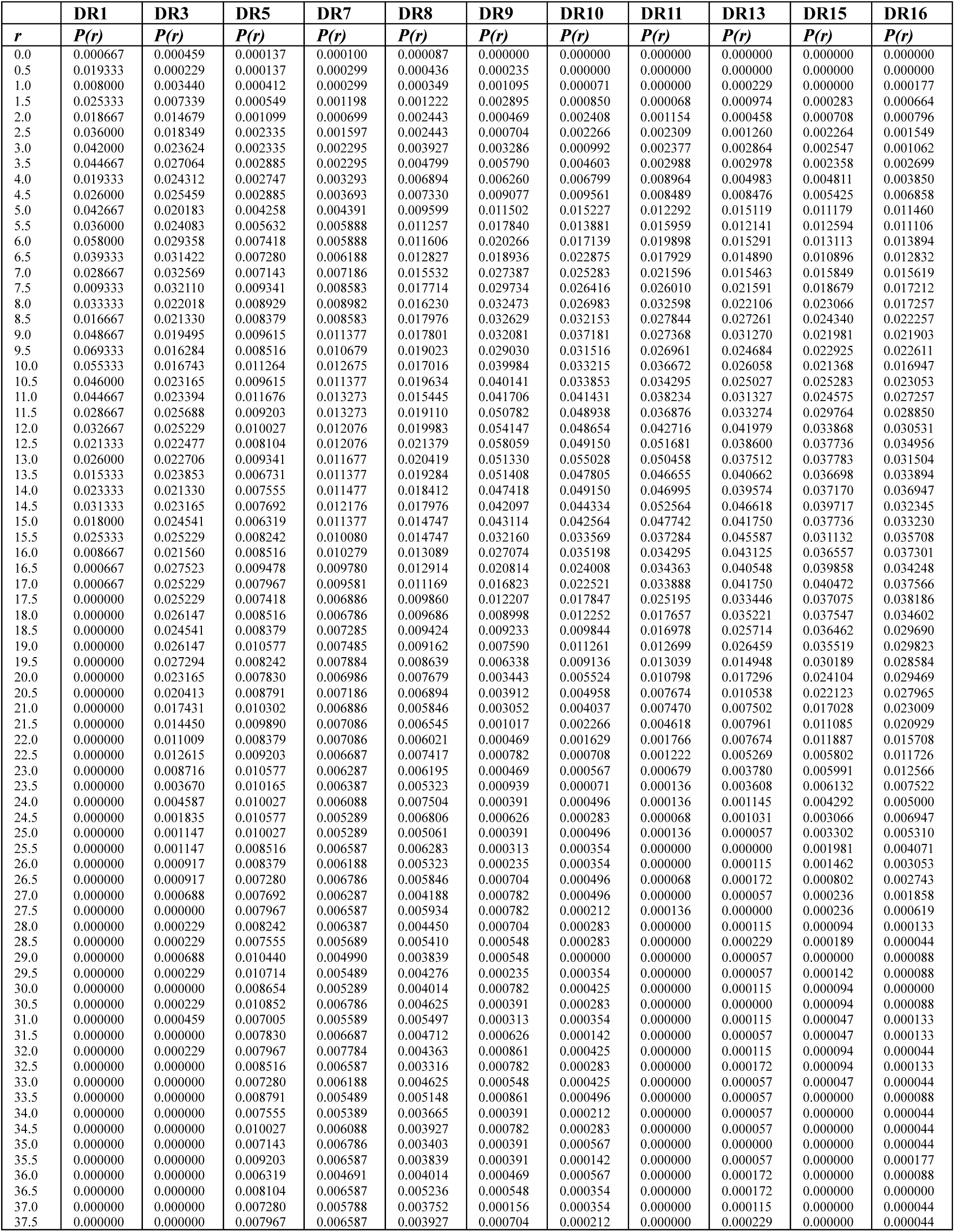
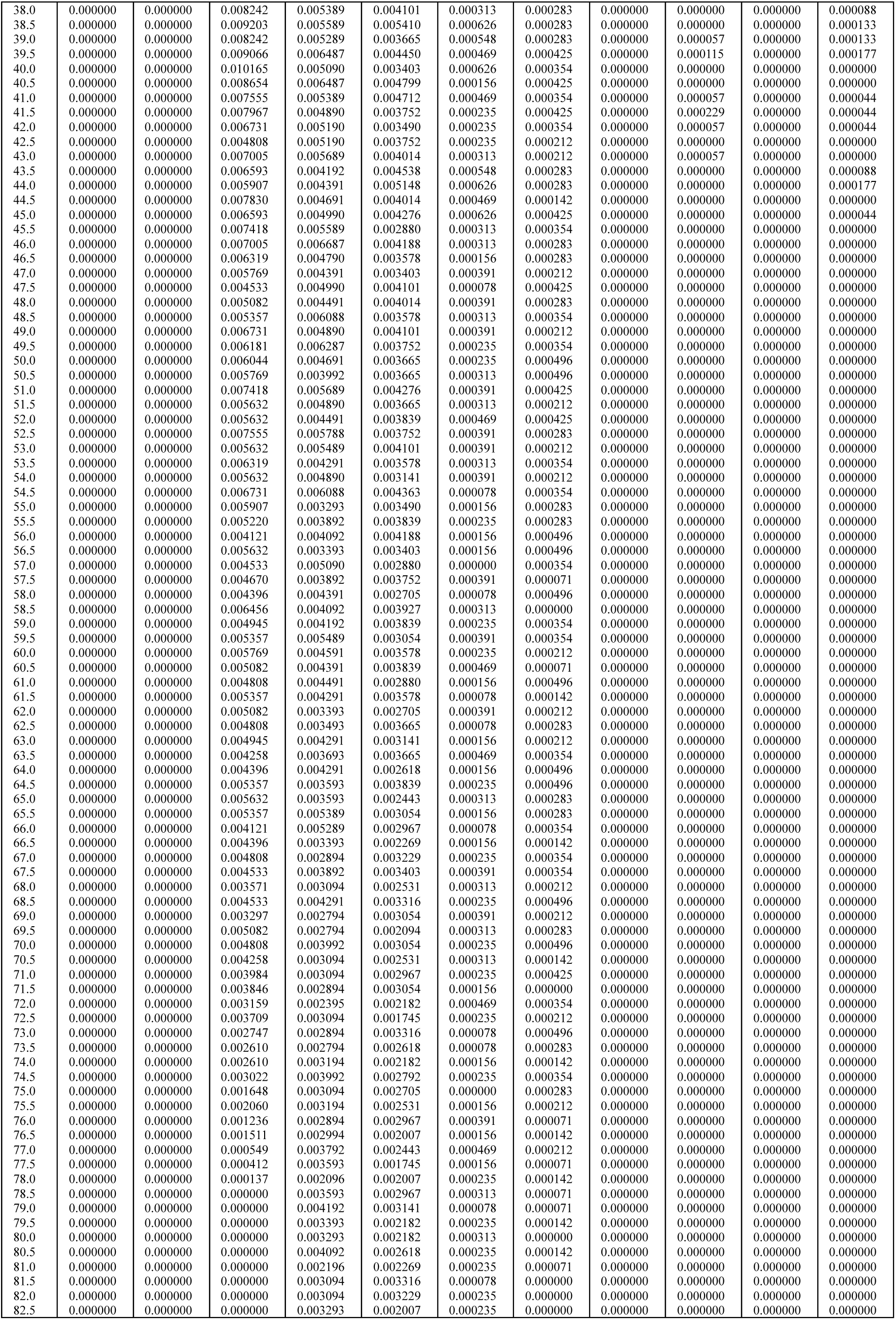
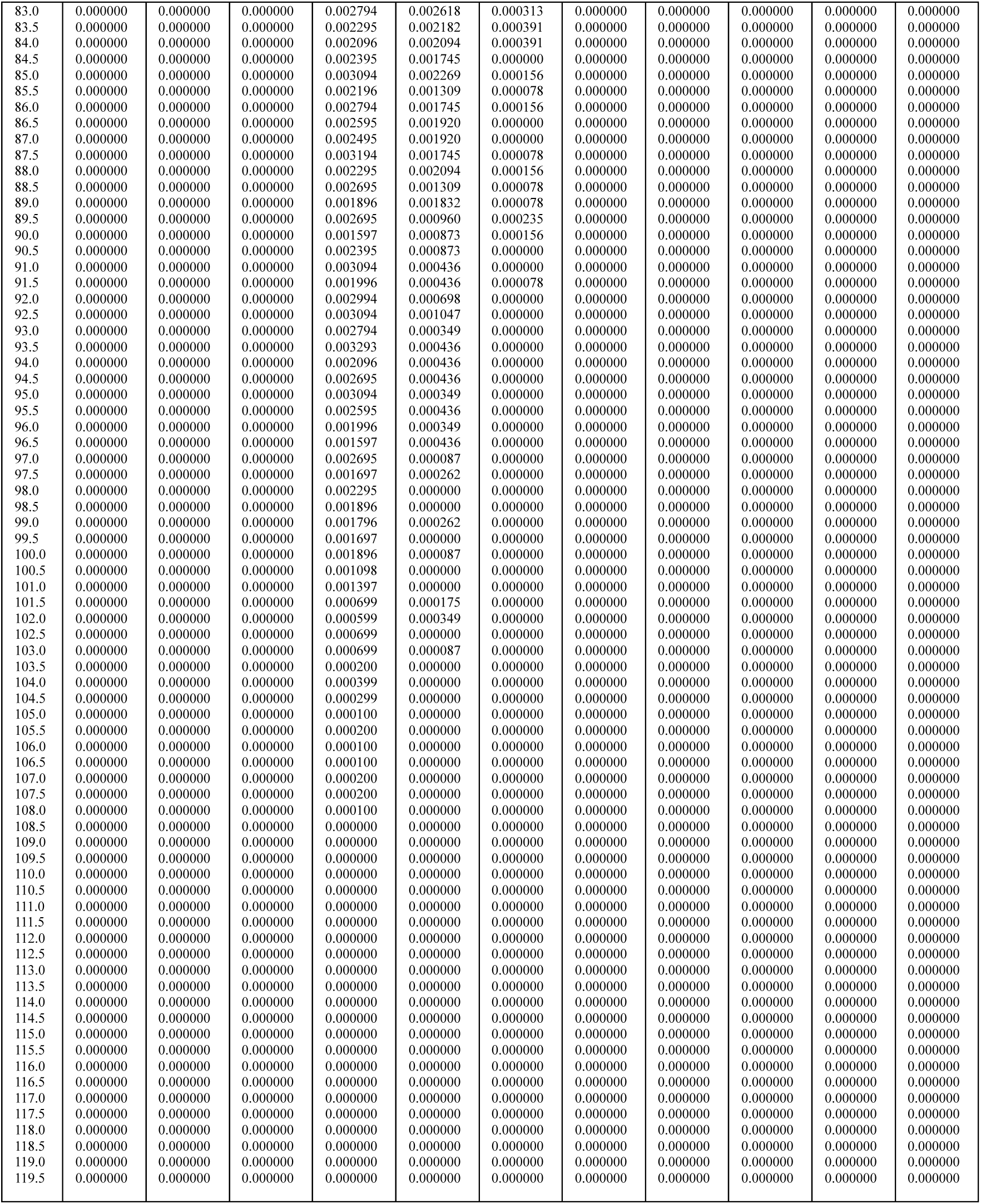
Raw data to Fig. 3d, Probability distributions *P (r)* of repeat atom distances *r* from the geometric centers of the repeats are shown for glycan-free repeats as predicted by AlphaFold3

**Figure 3.**
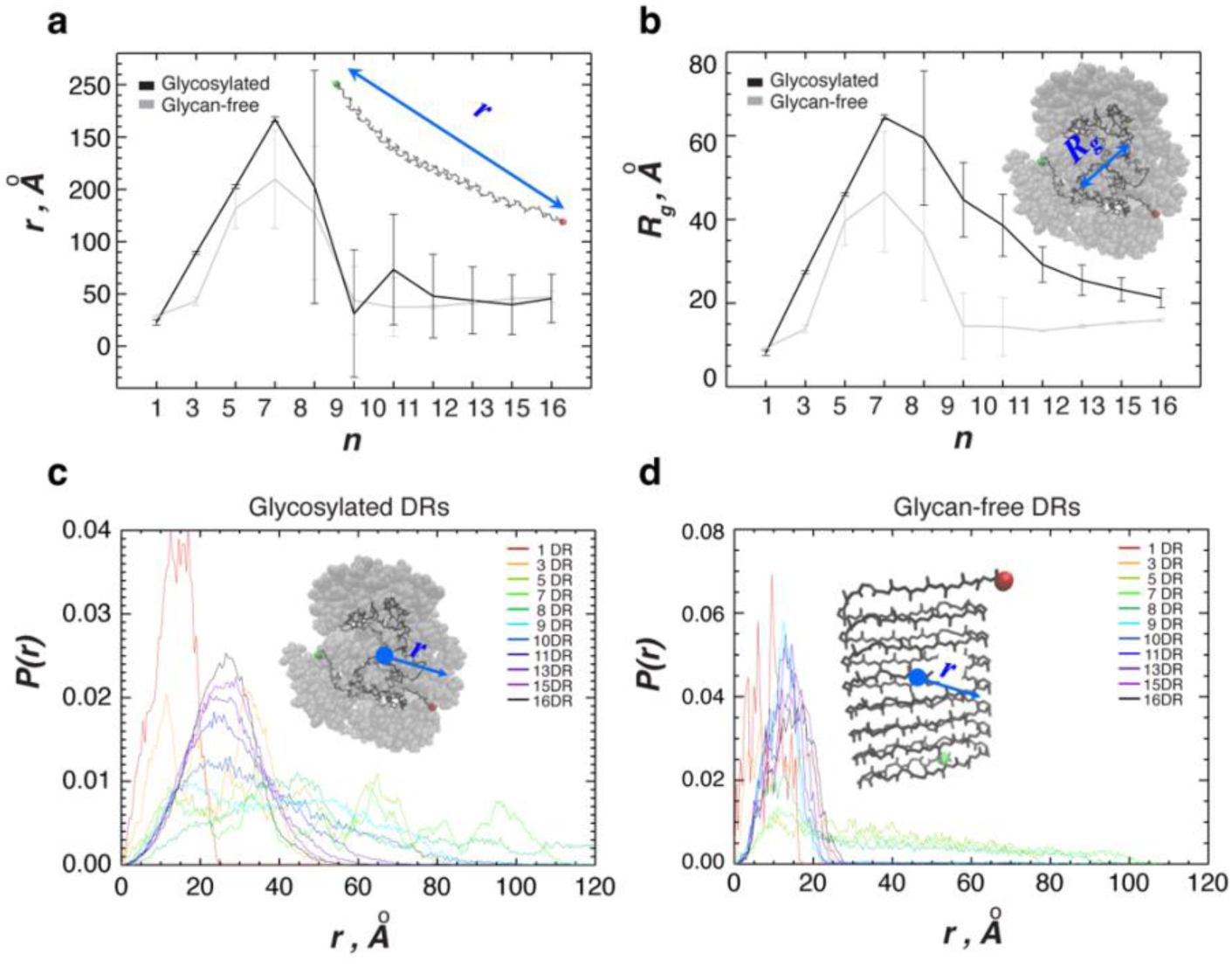
Conformational properties of DR repeats predicted by AlphaFold 3. (**a**) End-to-end distance, *r*, for DR repeats is shown as a function of the number of DRs, *n*. The end-to-end distance measures the separation between the first (in green) and last (in red) amino acids in the molecule. (**b**) The radius of gyration, *R_g_*, of DR repeats is shown as a function of the number of DRs, *n*. The radius of gyration measures the overall dimension of a molecule. In both panels, the data for glycosylated and glycan-free DRs are in black and grey, respectively. Vertical bars represent standard deviations for a given *n*. (**c** and **d**) Probability distributions *P*(*r*) of DR atom distances *r* from the geometric centers of the repeats are shown for glycosylated (**c**) and glycan-free (**d**) repeats. Different colors correspond to DR repeats with different *n*. All panels indicate that glycosylated DR repeats extend up to *n* = 7, beyond which the molecule collapses, which continues with the further increase in the number of repeats until *n* = 16. Glycan-free repeats demonstrate a similar extension-collapse transition, with the exception that long glycan-free repeats with *n* > 7 collapse into a β-sheet structure, which is notably more compact than that of the glycosylated repeats.

**Figure 4.**
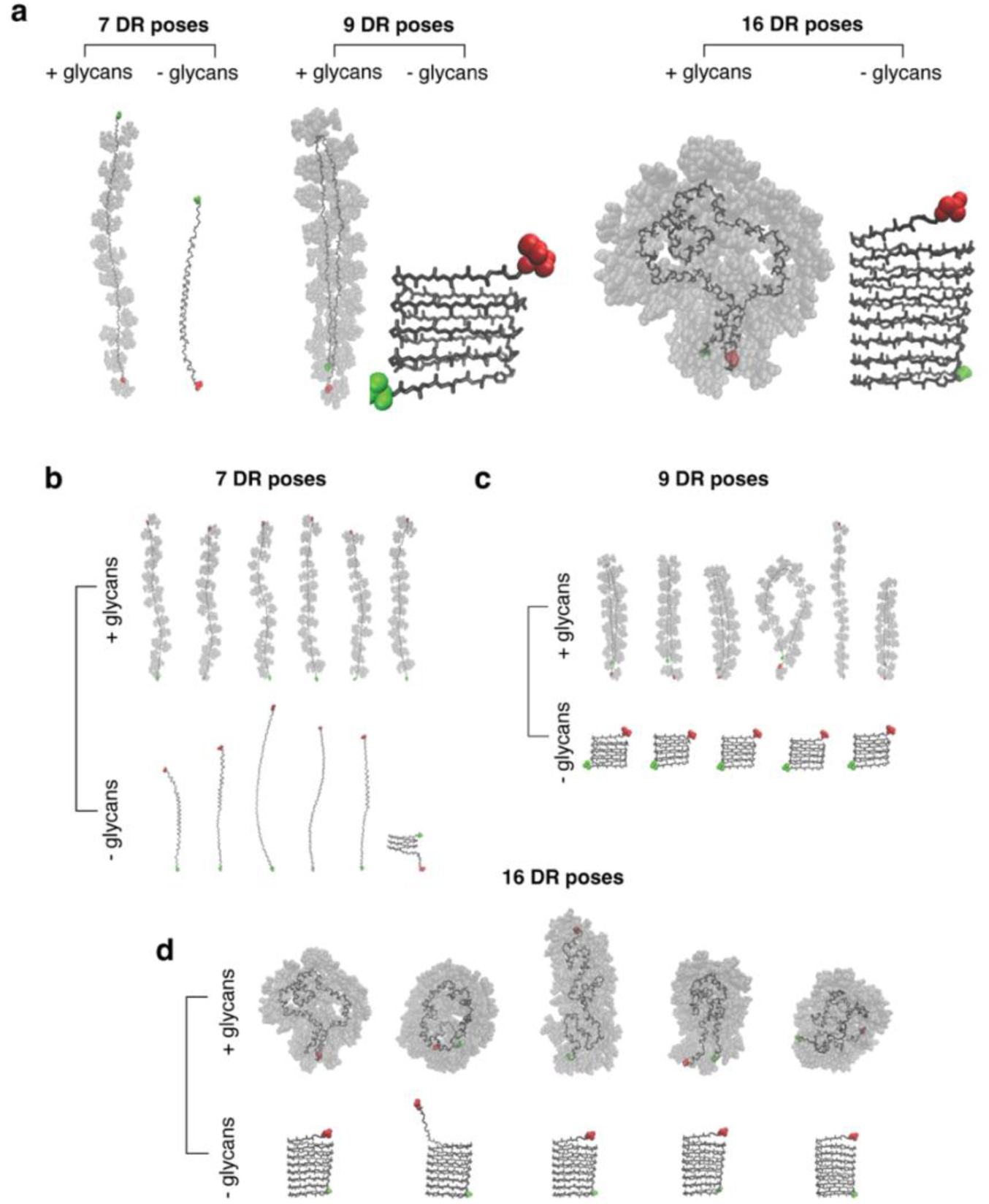
Predicted structures of DR repeats. (**a**) Representative structures of glycosylated (left) and glycan-free (right) DR repeats with lengths *n* = 7, 9, and 16 predicted by AlphaFold 3. (**b** to **d**) Distributions of structures of DR repeats predicted by AlphaFold 3. The first and last residues in each repeat are shown in green and red, respectively. Protein heavy atoms are shown in black, with glycans represented by translucent grey volume-filling spheres. Apart from some structural variability, these panels illustrate the extension-collapse transitions implicated in Fig. 3. Glycosylated DR repeats with *n* = 7 are extended, whereas the addition of two extra DRs forces the repeats to collapse. The extension from *n* = 9 to 16 results in further molecular compaction. Although the glycan-free DR repeats follow the same extension-collapse transition between *n* = 7 and 9, these molecules adopt beta-sheets for *n* = 9 and 16.

## Discussion

In this article, we investigated the role of PSGL-1’s extracellular (EC) and intracellular (IC) domains in HIV Env exclusion and found that both EC and IC are important for Env exclusion. In particular, EC and its DR domain play an essential role for PSGL-1-mediated Env exclusion. Sequential DR deletion mutagenesis further demonstrated that a minimum of 9 DRs is necessary for Env exclusion in our bioassays. We further performed molecular modeling of the DR repeats, and surprisingly, it revealed that there is a dramatic structural transition when the number of DRs extends beyond 7. PSGL-1 poses as a “rod-like” structure when the number of DRs is 7 or fewer. However, when *n* is 8 or more, the repeats collapse into a larger “coil-like” structure, while the DR repeat footprint projected on the virion surface grows almost three-fold, from 10Å at *n* = 7 to 27Å at *n* = 16. Importantly, this structural transition to a larger size is coincident with our experimental data showing that a minimum of 9 DRs is required for Env exclusion (**Fig. 2**). Based on these results and the structural information of HIV Env trimers ^22–24^, we proposed a model suggesting that PSGL-1-mediated Env exclusion likely involves Gag-mediated PSGL-1 targeting to the virion assembly site where DR-mediated spatial exclusion blocks Env incorporation (**Fig. 5**). The heavily glycosylated DRs are necessary for Env exclusion. However, the exact molecular details of how DRs facilitate Env exclusion remain to be determined. It is possible that there is a direct spatial competition between gp120 trimers and PSGL-1 on the plasma membrane when DRs collapse into the larger coil-like structure (**Fig. 5**). Although HIV virions carry only approximately 14 trimers per particle, Env trimers likely cluster (15 nm, the most frequent nearest-neighbor inter-spike spacing) and are not free to diffuse in the plane of the plasma membrane (23 nm, inter-spike spacing predicted from random distribution) ^24^. DR-collapsed PSGL-1 may have a volume large enough to affect inter-spike spacing, reducing the total numbers of gp120 trimer on the plasma membrane. Additionally, there may be inter-spike interaction mediated by the Env cytoplasmic tails ^24^, and PSGL-1 may also interfere with the Env cytoplasmic tail interactions through MA ^14^.

**Figure 5.**
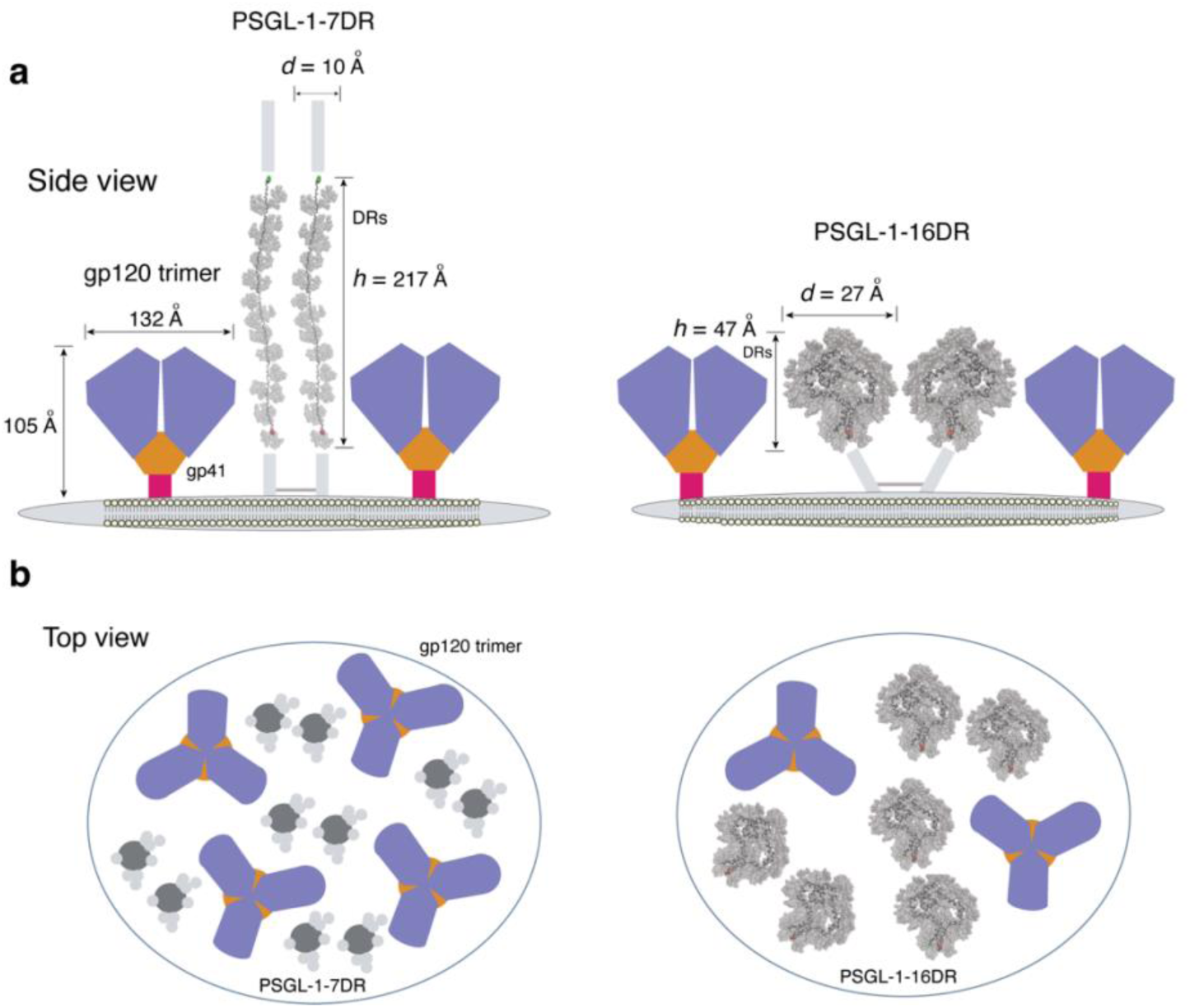
Hypothetical model by which virion incorporation of PSGL-1 inhibits HIV Env incorporation. (**a** and **b**) On the virion particle, PSGL-1 DR mutants can pose either as a “rod-like” structure when the number of DRs is 7 or fewer, or as a collapsed “coil-like” structure when *n* is 9 or more. Only the full-sized PSGL-1 or PSGL-1 mutants with 9 or more DRs spatially inhibit Env incorporation. The height *h* of DR repeats measured by their end-to-end distance (see Methods) decreases more than four-fold, from 217 Å at *n* = 7 to 47 Å at *n* = 16. Simultaneously, the dimension of the DR repeat footprint *d* grows almost three-fold, from 10 Å at *n* = 7 to 27 Å at *n* = 16. To compute *d*, we used the radius of gyration projected on the surface of the virion particle and considered all heavy atoms. The structural information of Env trimer is based on ^22,24^

Interestingly, the modeling results in **Fig. 3** indicate that the extension-collapse transition in DR repeats occurs regardless of DR glycosylation. However, the presence of glycans does affect the way in which the DR structures collapse. Indeed, the modeling in **Fig. 4** shows that long glycan-free DR repeats collapse into β-sheets, whereas glycosylated repeats collapse into disordered coil-like structures. Both extension-collapse transitions are likely driven by the competition between chain sterics, entropy, and the interactions within DRs. Our modeling results are corroborated by recent simulations of mucin fragments, which show that glycosylation increases protein dimensions and its rigidity as quantified by the persistence length ^25^.

PSGL-1 DR mutants with fewer (7 DRs or fewer) or no DRs lack the ability to exclude Env from virions. However, we also found that PSGL-1 mutants carrying a single DR have basal anti-HIV activity ^26^, suggesting that for PSGL-1, inhibiting virion attachment through “steric hindrance” is a dominant phenotype over its capacity to exclude Env through “spatial hindrance”.

## Methods

### Cells and cell culture

HEK293T cells were obtained from ATCC and cultured in 1x DMEM (Thermo Fisher Scientific) supplemented with 1x NEAA (GIBCO) 10% heat-inactivated FBS (Neuromics) and penicillin and streptomycin (Thermo Fisher Scientific). The cells were incubated at 37°C and 5% CO_2_.

### Plasmids and vectors

The infectious HIV-1 plasmid pNL4-3 was obtained from the NIH AIDS reagent program. The PSGL-1 mutants pcDNA3.1-PSGL-1, pcDNA3.1-PSGL-1-NT, pcDNA3.1-PSGL-1-CT, pcDNA3.1-PSGL-1-ΔDR(YH), pcDNA3.1-PSGL-1-1DR #16, pcDNA3.1-PSGL-1-3DR, pcDNA3.1-PSGL-1-7DR, pcDNA3.1-PSGL-1-9DR, pcDNA3.1-PSGL-1-11DR, pcDNA3.1-PSGL-1-13DR, and pcDNA3.1-PSGL-1-15DR were synthesized by Twist Bioscience. pCMV3-PSGL-1 and pCMV3-empty vectors were obtained from Sinobiological. The plasmid pPSGL-1ΔCT was kindly provided by Dr. Ono’s Lab ^14^.

### Transfection and virion production and purification

HEK293T cells were co-transfected with pNL4-3 (1µg) and individual PSGL-1 mutants (500 ng) or pCMV plasmid as empty vector as previously described ^17^. Supernatants were collected after 48 hours, and viral particles were purified using a low-speed virus concentration kit (Virongy).

### SDS-PAGE, Western blotting, and antibodies

Purified virions were resuspended in 1x lysis buffer (Thermo Fisher Scientific) containing 1x Protease Inhibitor Cocktail (Thermo Fisher Scientific), vortexed, and boiled at 100°C for 5 min. The samples were then centrifuged at 13,000 rpm for 5 min. Proteins were separated using Bis-Tris Mini Protein Gel (4-12%) (Thermo Fisher Scientific) and then transferred to a nitrocellulose membrane (Thermo Fisher Scientific). The blots were blocked with 5% skim milk (Difco) in 1x TBST, then incubated overnight at 4°C with primary antibodies as follows: anti-HIV-1 gp41 antibody (Chessie8) (1:1000) (NIH AIDS Reagent Program); anti-PSGL-1 (KPL-1) (1:1000) (BioLegend); anti-HIV-1 p24 antibody (183-H12-5C) (NIH AIDS Reagent Program) (1:1000) in 1x TBST with 2.5% skim milk. Anti-mouse IgG, HRP-linked Antibody (1:1000) (Cell Signaling) was used as the secondary antibody. The blots were developed using Chemiluminescent Substrate, ECL (Thermo Fisher Scientific), and images were acquired using a Bio-Rad ChemiDoc Imaging system and analyzed with Bio-Rad Image Lab software.

### Software and modeling methods

AlphaFold 3 ^20^ was used to predict the structures of PSGL-1 DR repeats with and without glycans. PSGL-1 is known to have O-glycans with the sialyl Lewis x antigen ^27^. Because this glycan is not available in AlphaFold 3, it was approximated in our modeling using a similarly branched glycan composed of 2-acetamido-2-deoxy-beta-D-glucopyranose, alpha-L-fucopyranose, and alpha-D-mannopyranose. Both glycans have similar sizes and net charges of −2 and 0, respectively. Although we used the glycans with zero net charge, one may expect that electrostatic repulsion between O-glycans is diminished at physiological salt concentrations. For each DR repeat, 20 unique AlphaFold runs were performed, with 5 models output per run. The highest-scoring models from each run were used for analysis. This procedure was repeated for both glycosylated and glycan-free DRs. All modeling data are presented as averages across the best models in 20 AlphaFold runs.

In modeling the PSGL-1 monomeric chain, we also assumed that the PSGL-1 dimeric state does not affect the conformational properties of its monomeric chains. This assumption appears reasonable given the highly polar character of glycans.

## Data analysis

Statistical analyses were performed with GraphPad Prism 10.0 (GraphPad Software). Two-way ANOVA was used to test the significance of protein expression among the samples. Applied statistical analyses are indicated in the figure legends. Data are presented as mean ± SEM of three replicates unless stated otherwise in the figure legend.

## Data availability

All data are available from the manuscript and the related Extended Data.

## Acknowledgments

The authors wish to thank the NIH AIDS Reagent Program for reagents. This work was supported by Public Health Service grant R01AI148012 (to Y.W.) and R56AI183995 (to Y.W., S.J., and D.K.K). Research in the Freed laboratory is supported by the Intramural Research Program of the Center for Cancer Research, National Cancer Institute, NIH and the Intramural AIDS Targeted Antiviral Program.

## Author contributions

Y.W. conceived the project. Y.W. and E.O.F. performed experimental data analysis. D.K. and M.S.J. performed and supervised molecular modeling. B.M.D. and C.L. performed molecular modeling. S.T., Y. H., A.H., and A.A.W. performed DNA mutagenesis and virology experiments and data analysis. Y.W. wrote the manuscript draft. Y.W. and D.K. supervised all work. Y.W. and D.K. prepared the manuscript with input from all authors.

